# Extensive genetic diversity of bat-borne polyomaviruses reveals inter-family host-switching events

**DOI:** 10.1101/627158

**Authors:** Zhizhou Tan, Gabriel Gonzalez, Jinliang Sheng, Jianmin Wu, Fuqiang Zhang, Lin Xu, Peisheng Zhang, Aiwei Zhu, Yonggang Qu, Changchun Tu, Michael J. Carr, Biao He

**Affiliations:** Key Laboratory of Jilin Province for Zoonosis Prevention and Control, Institute of Military Veterinary Medicine, Academy of Military Medical Sciences, Changchun, Jilin Province, China; National Advanced Computing Collaboratory, National Center for High Technology, San Jose, Costa Rica; College of Animal Science and Technology, Shihezi University, Shihezi, Xinjiang Uyghur Autonomous Region, China; Guangxi Key Laboratory of Veterinary Biotechnology, Guangxi Veterinary Research Institute, Nanning, Guangxi Province, China; Center for Disease Control and Prevention of Southern Theater Command, Kunming, Yunnan Province, China; Jiangsu Co-innovation Center for Prevention and Control of Important Animal Infectious Diseases and Zoonosis, Yangzhou, Jiangsu Province, China; National Virus Reference Laboratory, School of Medicine, University College Dublin, Dublin 4, Ireland

## Abstract

Polyomaviruses (PyVs) are small, double-stranded DNA tumor viruses carried by diverse vertebrates. PyVs have previously been considered highly host restricted in mammalian hosts, with host-switching events thought rare or nonexistent. Prior investigations have revealed short-range host-switching events of PyVs in two different African bat species within the horseshoe bat genus *Rhinolophus*. Herein, we have conducted a systematic investigation of PyVs in 1,083 archived bat samples collected from five provinces across China, and identified 192 PyVs from 186 bats from 15 host species within 6 families (Rhinolophidae, Vespertilionidae, Hipposideridae, Emballonuridae, Miniopteridae and Pteropodidae) representing 28 newly-described PyVs, indicative of extensive genetic diversity of bat PyVs. Surprisingly, two PyVs were identified in multiple bat species from different families, and another PyV clustered phylogenetically with PyVs carried by bats from a different host family, indicative of three inter-family PyV host-switching events. The time to most recent common ancestor (tMRCA) of the three events was estimated at 0.02-11.6 million years ago (MYA), which is inconsistent with the estimated tMRCA of their respective bat hosts (36.3-66.7 MYA), and is most parsimoniously explained by host-switching events. PyVs identified from geographically separated Chinese horseshoe bat species in the present study showed close genetic identities, and clustered with each other and with PyVs from African horseshoe bats, allowing assessment of the effects of positive selection in VP1 within the horseshoe bat family Rhinolophidae. Correlation analysis indicated that co-evolution with their hosts contributed much more to evolutionary divergence of PyV than geographic distance. In conclusion, our findings provide the first evidence of inter-family host-switching events of PyV in mammals and challenge the prevailing evolutionary paradigm for strict host restriction of mammalian PyVs.

**Author summary:** Since the discovery of murine polyomavirus in the 1950s, polyomaviruses (PyVs) have been considered both genetically stable and highly host-restricted in their mammalian hosts. In this study, we have identified multiple cases of host-switching events of PyVs by large scale surveillance in diverse bat species collected in China. These host-switching events occurred between bat families living in the same colony, indicating that a large population with frequent contacts between different bat species may represent an ecological niche facilitating PyV host-switching. The cases studied involved members of bats from several families, including horseshoe bats, which were previously found to harbor a number of highly virulent viruses to both humans and domestic animals. Our findings have provided evidence that even highly host-specific DNA viruses can transmit between bats of different species and indicate an increased propensity for spillover events involving horseshoe bats. We propose an evolutionary scheme for bat-borne PyVs in which intra-host divergence and host-switching has generated the diverse PyVs in present day bats. This scheme provides a useful model to study the evolution of PyVs in other hosts and, potentially, the modeling of bat zoonoses and the transmission of other DNA viruses in other mammals, including humans.

## Introduction

Bats (order Chiroptera) account for more than 20% of mammalian species with > 1,100 known species in > 200 genera worldwide [1]. These diverse mammals harbor numerous highly virulent viral pathogens, including rabies virus [2], Marburg virus [3], Ebola virus [4], Nipah virus [5], severe acute respiratory syndrome (SARS)-related coronavirus [6-8] and the newly identified swine acute diarrhea syndrome coronavirus [9]. Spillover of viruses from bats to humans or domestic animals can give rise to outbreaks of emerging infectious diseases [10].

Polyomaviruses (PyVs) are non-enveloped icosahedral DNA viruses containing a circular and highly stable double-stranded DNA genome [11, 12]. PyVs can induce neoplastic transformation in cell culture [12], and are associated with malignancy in humans [13, 14] and raccoons [15]. Currently, 98 species carried by various mammals, birds and fish are recognized within the family *Polyomaviridae* by the International Committee on Taxonomy of Viruses (ICTV) and classified into four separate genera (*Alpha-, Beta-, Gamma-* and *Deltapolyomavirus*) with nine floating species [16]. Recent identification of diverse PyV-like sequences in different fish species [17] and invertebrates (including arachnids and insects) has revealed a deep co-evolutionary history of PyVs with their metazoan hosts [17]. PyVs are highly host-specific in different mammalian species, and long-range host jumps leading to productive infection and transmission within mammalian genera are considered rare or nonexistent [17, 18]. Recently, an intra-host divergence model with contributions from viral lineage duplication, recombination and, potentially, host-switching at evolutionary timescales has been proposed to explain the extant diversity based on the available genetic data [17].

Currently, 15 alphapolyomaviruses and 10 betapolyomaviruses have been formally recognized in insectivorous and frugivorous bat species within the Americas, Africa, Southeast Asia (Indonesia) and the Pacific (New Zealand) [19-26]. These studies identified significant diversity and high positive rates (10-20 %) using PCR-based approaches [27]. PyVs with complete or partial genomes has also been identified by metagenomic analyses in bats collected in China and recently from Saudi Arabia [28-30]. Investigation of PyVs in African insectivorous and frugivorous bats [25, 26] has revealed evidence for short-range host-switching events of PyVs in horseshoe bats (family Rhinolophidae, genus *Rhinolophus*) following identification of nearly identical viruses (99.9% at the genomic level) in *Rh. blasii* and *Rh. simulator* [26].

In the present study, we have characterized the diversity and genetic relationships of novel PyVs from a large number of insectivorous and frugivorous bat samples collected in diverse ecological environments in China, resulting in the identification of inter-family host-switching events in these bat-borne PyVs. The findings challenge the prevailing paradigm of intra-host PyV diversification in their mammalian hosts and extend the knowledge of PyV genetic diversity, evolution, host restriction and host-switching.

## Results

### Sampling information and detection of polyomaviruses

In 2015-2016, a total of 1,083 bats belonging to 20 species in 6 families (Rhinolophidae, Vespertilionidae, Hipposideridae, Emballonuridae, Miniopteridae and Pteropodidae) were collected from 29 different colonies (colonies 1-29 in Fig 1 and S1 Table) from five provinces in southwest (Yunnan and Guangxi), southeast (Fujian and Zhejiang) and northwest (Xinjiang) provinces in China. The bats were identified morphologically by field-trained experts during sampling and confirmed by partial mitochondrial *cytochrome b* (*cytb*) gene sequencing [31] of representative individuals in every colony (S1 Table). The bats were insectivorous except for one fruit bat species *Rousettus leschenaultii* in the family Pteropodidae, and were captured in diverse habitats, including natural and artificial caves, orchards and human-made structures (S1 Table). Archived tissues from individual bats had previously been sampled for zoonotic virus surveillance, including for group A rotaviruses (RVA) [32] and hantaviruses [33].

**Fig 1.**
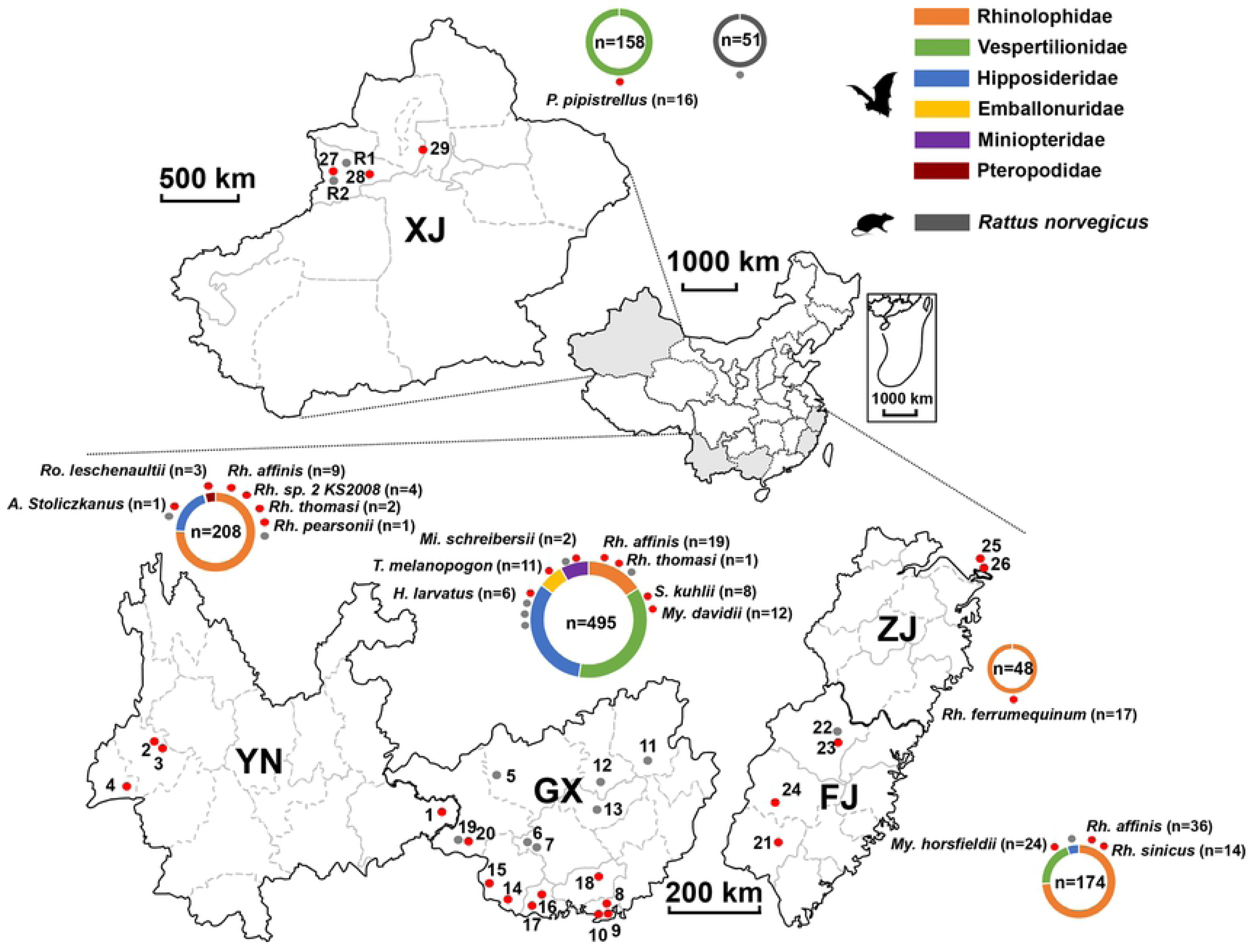
Sampling locations and species composition of animals collected in different provinces. The filled circles represent the locations of sampled colonies encoded by numbers (see details in S1 File) with the red ones representing PyV-positive colonies or mammalian species and the grey ones representing the PyV-negative colonies or mammalian species (species names of PyV negative bats are not shown). The composition of collected bats or rodents in each province is presented by the colored rings with total numbers in the rings. The color region in the rings represents each positive bat families or rodent species as indicated in the top right-hand corner. Every filled circle around the colored rings represents one mammalian species. The numbers of PyV positive bats in each species are shown in parentheses. Abbreviations of provinces: XJ, Xinjiang Uyghur Autonomous Region; YN, Yunnan Province; GX, Guangxi Zhuang Autonomous Region; FJ, Fujian Province; ZJ, Zhejiang Province. Abbreviations of bat genera: *H*., *Hipposideros; A*., *Aselliscus; Rh*., *Rhinolophus; S*., *Scotophilus; Mi*., *Miniopterus; My*., *Myotis; T*., *Taphozous; Ro*., *Rousettus*.

As part of the regional viral surveillance of a bat-borne zoonosis program, the lungs and rectal tissue of 208 bats of 9 bat species from colonies 1-4 (S1 Table) collected in Yunnan province were pooled and subjected to viral metagenomic analysis, as previously described [34]. Nucleic acids were amplified by sequence-independent single primer amplification prior to Illumina sequencing. A total of 1.76×10^7^ sequence reads with an average length of 125 nucleotides (nt) were obtained, of which 694 PyV-related reads with average length of 167 nt were identified from 38,468 virus-related sequences. Preliminary analyses of these PyV sequences indicated a high level of diversity with previously unrecognized members from both *Alpha*- and *Betapolyomavirus* genera.

To determine the prevalence of PyVs in bats collected not only from Yunnan but also from other provinces in China, a broad-spectrum PyV nested PCR targeting the VP1 gene of known members within the *Alpha-* and *Betapolyomavirus* genera [27] was performed using genomic DNAs extracted from archived rectal tissues of all 1,083 bats. This yielded 186 (17.2%) PyV positive samples from 15 bat species within 6 families. These PyV positive bats were from 21 of 29 bat colonies in the survey area. As shown in Fig 1 and Table 1, all five investigated provinces had PyV positive bats with detection rates of 9.6% (20/208) in Yunnan, 11.9% (59/495) in Guangxi, 10.1% (16/158) in Xinjiang, 42.5% (74/174) in Fujian and 35.4% (17/48) in Zhejiang. With respect to the bat families (Fig 1 and Table 1), 103 from 6 species of Rhinolophidae, 60 from 4 species of Vespertilionidae, 7 from 2 species of Hipposideridae, and 11, 2 and 3 respectively from 1 species each of Emballonuridae, Miniopteridae and Pteropodidae were PyV positive. At the species level, *Rh. affinis*, in the Rhinolophidae, widely distributed in southern China, yielded the highest number of positive bats (n=64), while *Myotis horsfieldii* in the Vespertilionidae collected in Fujian, showed the highest positivity rate: 66.6% (24/36).

**Table 1.**
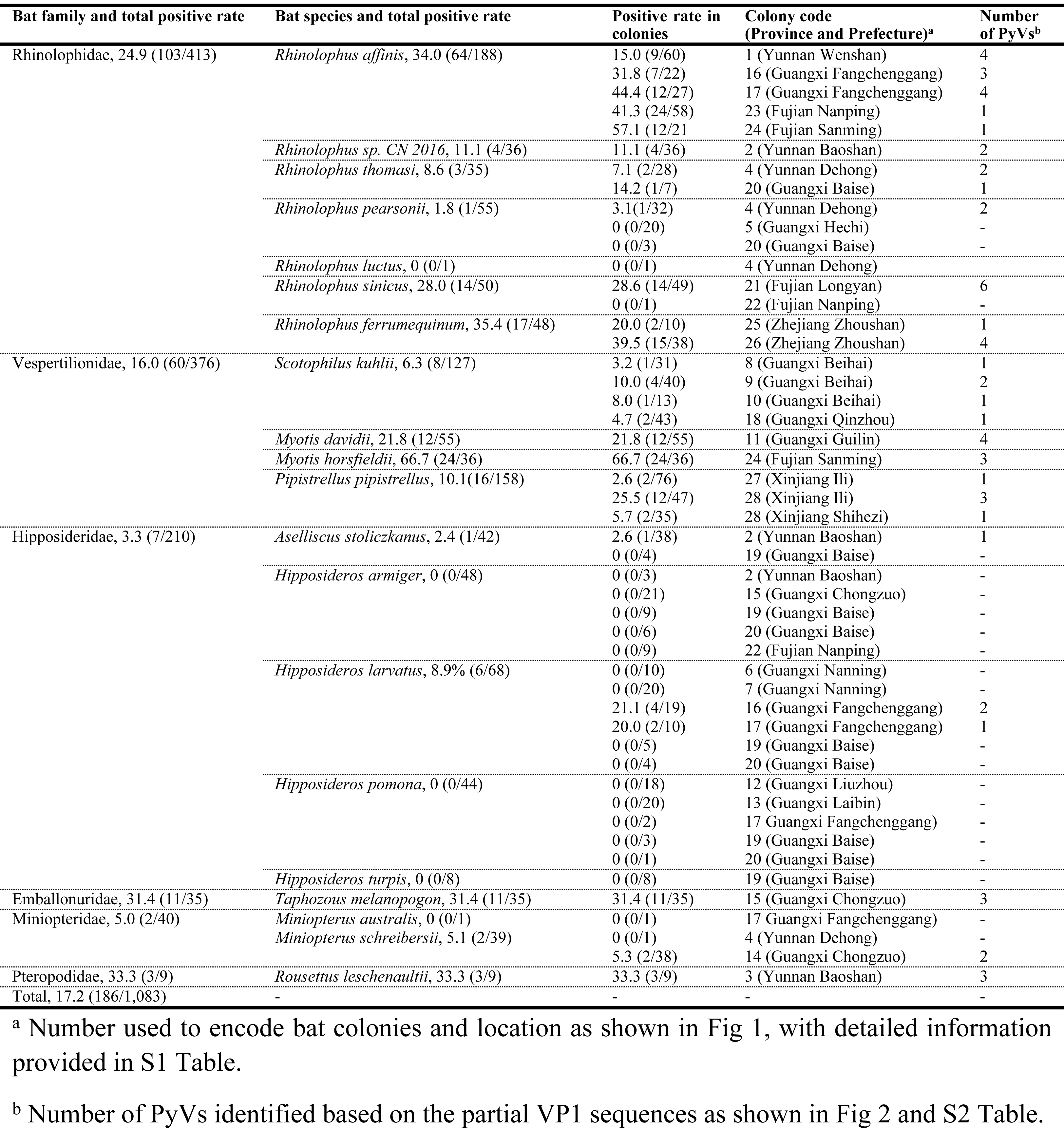
Sampling of bats and positive rate (%) of PyVs in Chinese provinces.

### Extensive genetic diversity of polyomaviruses among bats

VP1 genes of PyV were amplified using broad-spectrum primers from 186 positive samples and then cloned into vectors. Four clones derived from each amplicon were sequenced to detect potential co-infection of multiple PyVs in each positive sample. As a result, 180 of 186 bats were confirmed to harbor one PyV, while the remaining 6 had a co-infection with two distinct PyVs (detailed in S1 Table). A total of 192 non-redundant partial VP1 sequences were obtained and phylogenetically compared with VP1 sequences of known PyVs. Results showed that the 192 VP1 sequences had extensive diversity and could be classified into 44 clusters within the *Alpha-* and *Betapolyomavirus* genera (Fig 2 and S1 Fig), with 46.6-90.1% nt identities between clusters and 95.1-100% within clusters. Of these 44 PyVs clusters, 42 showed exclusive host specificity within a single bat species while, surprisingly, the remaining 2 clusters included viruses detected in two bat species within different families (*Rh. affinis* and *Hipposideros larvatus; Rh. affinis* and *Myotis horsfieldii*, respectively). This finding provided preliminary evidence of inter-family transmission of PyVs (Fig 2 and S1 Fig). The majority (n= 191) of these partial VP1 shared less than 90% nt identity with the closest known PyVs. Interestingly, one VP1 sequence (marked by cluster 10 in S1 Fig), named here as Pipistrellus pipistrellus polyomavirus XJPp02 (strain PyV10-YXAC25) shared 98% nt identity with Rattus norvegicus polyomavirus 1, previously identified in brown rats in Germany [35].

**Fig 2.**
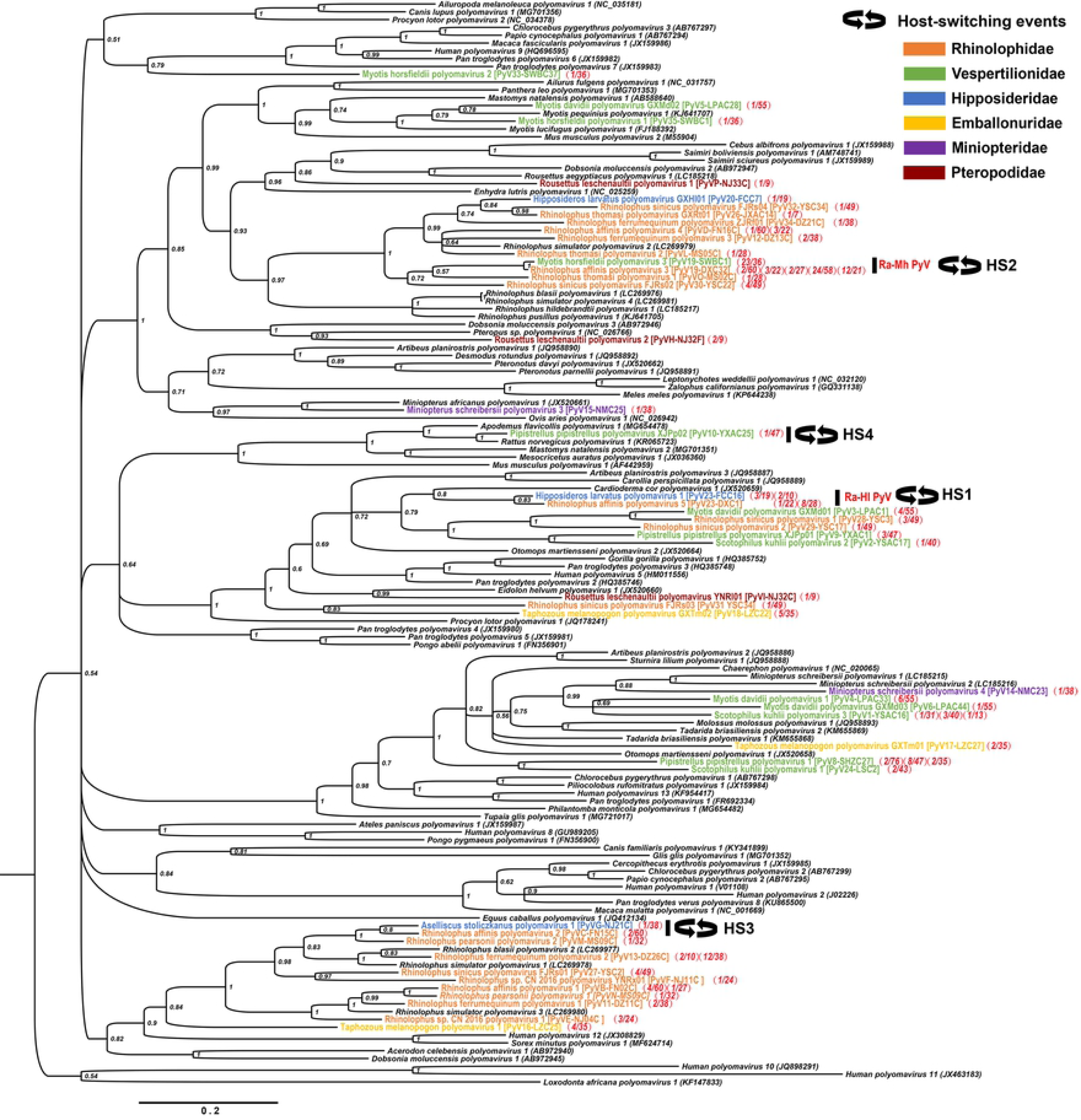
Bayesian phylogenetic tree of partial VP1 nucleotide sequences from PyV positive bats. The tree was constructed with 46 partial VP1 sequences (non-black-colored) of 44 new PyVs in present study, and 93 PyV references (including 38 reference bat PyV species) in *Alpha-* and *Betapolyomavirus* genera available in GenBank. The names of putative novel PyV strains are differentiated using colors according to the Chiropteran family of bat hosts following the legend in the top right-hand corner. The PyV positive rate of each putative PyV species from different colonies is shown in red in parentheses next to the tip names. Each parenthesis represents one colony. Host-switching events are marked in the clusters showing the presence of closely related PyVs with different bat host species (HS1-HS4). The posterior probability supporting the branching is shown next to the branch. Scale: 0.2 nucleotide mutations per site.

To further characterize the genetic diversity of the PyVs in these bats, inverse PCR based on the partial VP1 sequences was employed to amplify the remaining genomes. A total of 42 full genomes were obtained from 40 individual bats. As shown in Table 2, the genome sizes of the novel bat-borne PyV were 4,869–5,529 bp with 39.49–44.69% G+C content. All PyV genomes displayed an archetypal genomic organization, including an early region encoding two nonstructural proteins [large tumor antigen (LTAg), and short tumor antigen (STAg)], and a late region encoding three capsid proteins (VP1, VP2 and VP3) on opposite strands. Early and late regions were separated by a non-coding control region harboring regulatory elements. For all samples from which PyVs genomes were obtained, a ∼1,800 bp fragment of the *cytb* gene was amplified to corroborate the bat host species (Table 2 and S2 Fig).

**Table 2.**
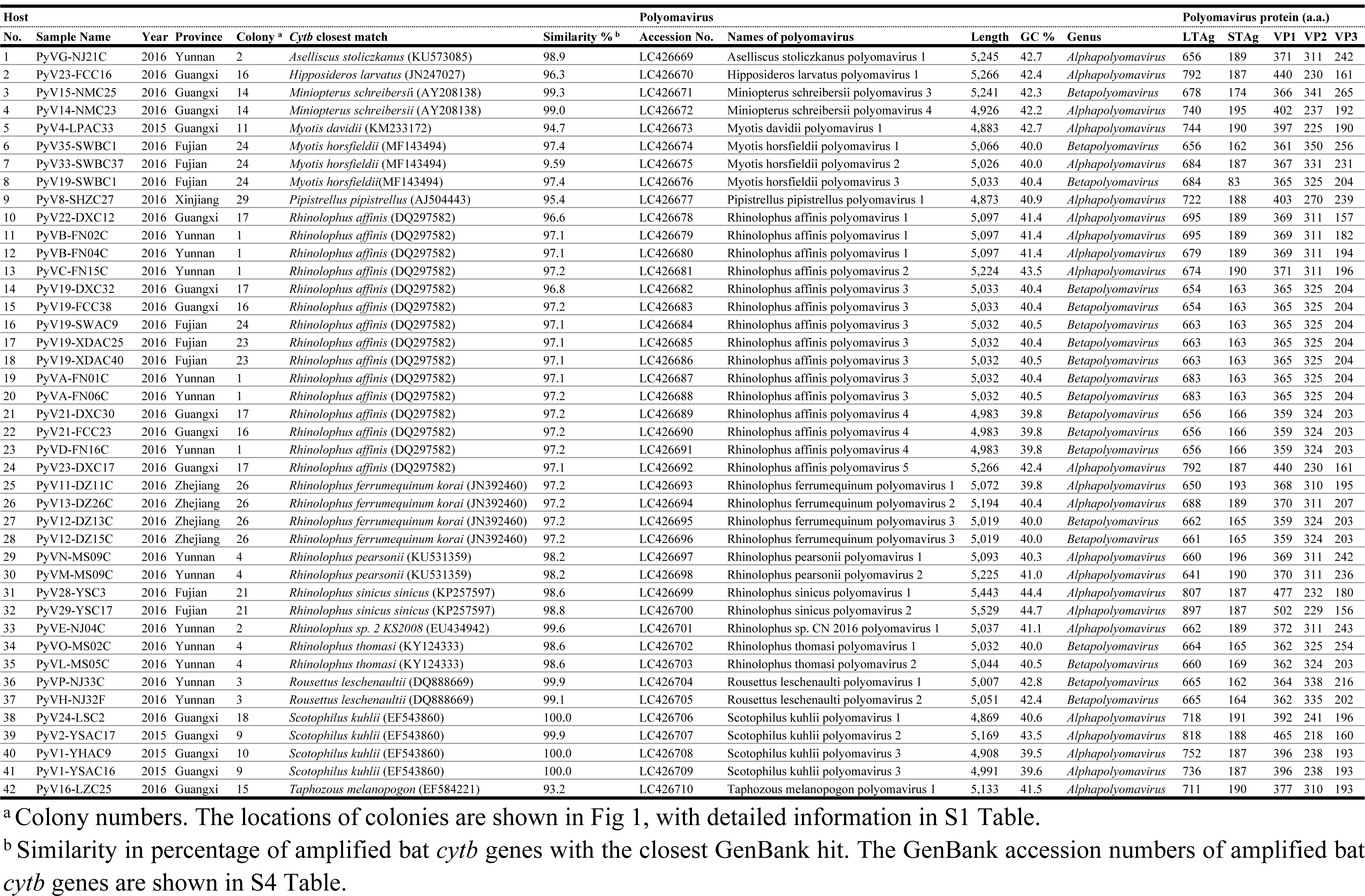
Novel bat PyV genomes identified in Chinese bat hosts.

The PyV strains sharing ≥ 97% nt identity within the LTAg coding sequence were defined as the same PyV, while those sharing ≤ 90% identities were defined as different PyV [36]. Consequently, as shown in Table 2 and Fig 3, the 42 PyV strains were identified as 28 novel bat-borne PyVs based on genomic identities. Of these novel bat-borne PyVs, 26 were identified in a single bat species, indicative of a high degree of host specificity. They included one PyV identified in a cryptic bat species (provisionally named *Rhinolophus sp. CN2016*) the permanent identification of which awaits formal binomial classification of the host species as the *cytb* gene showed only 90% nt identity with that of known bat species [37]. The PyVs named in the present study were based on the binomial names of their host bat species and the chronological order of discovery, as per ICTV guidelines. For example, Aselliscus stoliczkanus polyomavirus 1 was the first PyV discovered in *Aselliscus stoliczkanus* bats, and Rhinolophus affinis polyomavirus 2 was the second PyV discovered in *Rh. affinis* bats. Interestingly, two remaining PyVs were identified in more than one bat species. In detail, a PyV, herein provisionally named as Rhinolophus affinis-Hipposideros larvatus polyomavirus (Ra-Hl PyV), was identified in both *Rh. affinis* and *H. larvatus* bats. Similarly, another PyV named as Rhinolophus affinis-Myotis horsfieldii polyomavirus (Ra-Mh PyV) was identified in *Rh. affinis* and *My. horsfieldii* bats. For clarity and according to the naming tradition of PyV, the Ra-Hl PyV identified in *Rh. affinis* was named Rhinolophus affinis polyomavirus 5 (the fifth PyV discovered in *Rh. affinis*), and the Ra-Hl PyV identified in *H. larvatus* was named Hipposideros larvatus polyomavirus 1. Similarly, the Ra-Mh PyV identified in *Rh. affinis* was named Rhinolophus affinis polyomavirus 3, and the Ra-Mh PyV identified in *My. horsfieldii* named Myotis horsfieldii polyomavirus 3.

**Fig 3.**
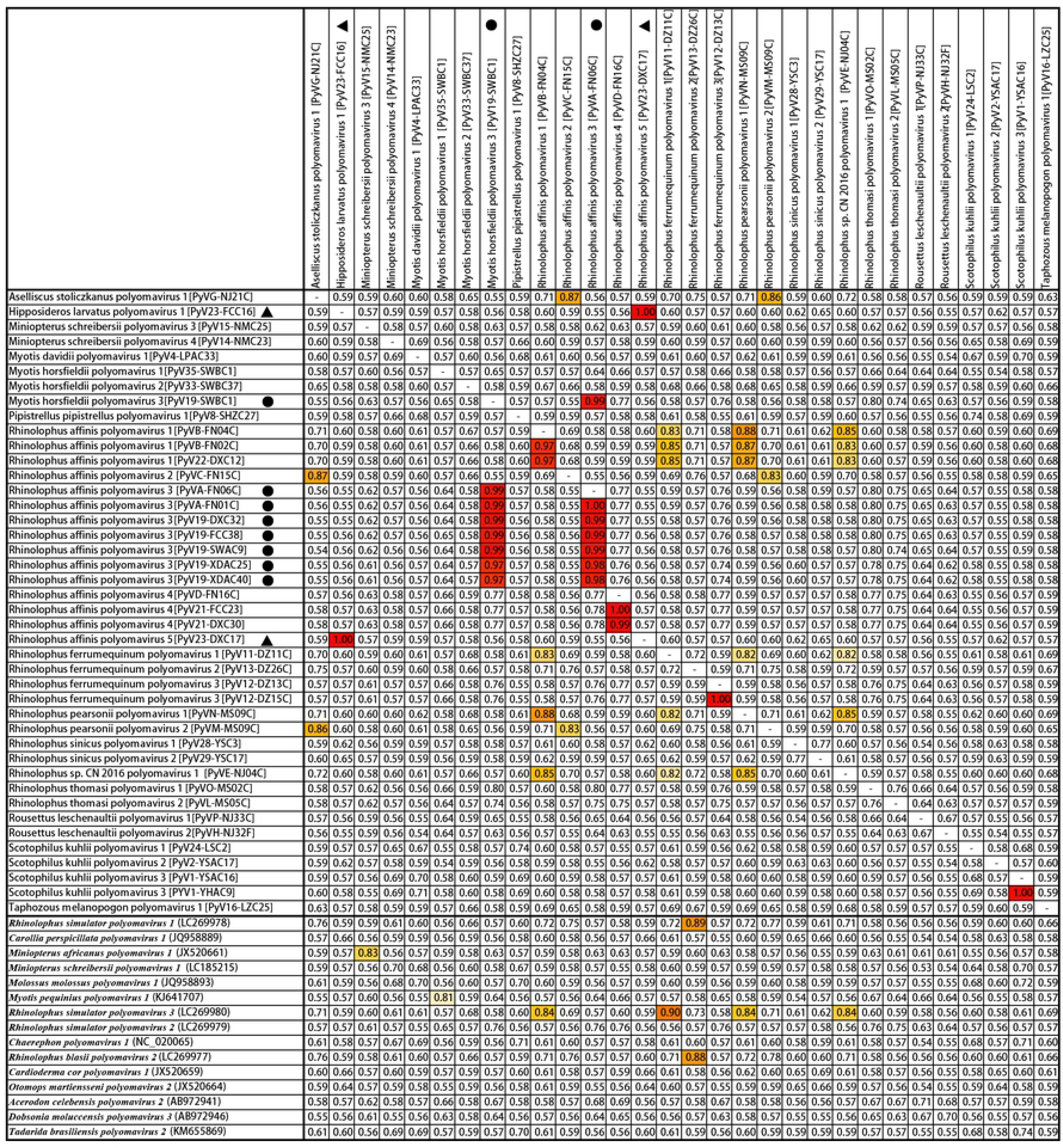
Pairwise identity comparison of the LTAg genes of bat-borne PyVs identified in the present study. The LTAg coding sequences of 42 bat-borne PyV strains identified in the present study and 15 PyV references (row names) are compared with those of 30 PyVs representing 28 newly proposed PyVs (column names). Cells are colored according to their nt sequence identity in red (> 0.90), orange (0.85-0.90), yellow tones (0.80-0.84) and white (< 0.80). Yellow and light yellow: nt identity 0.80-0.84. Rhinolophus affinis-Hipposideros larvatus polyomavirus (Ra-Hl PyV) are identified by triangles and Rhinolophus affinis-Myotis horsfieldii polyomavirus (Ra-Mh PyV) by circles.

Among the newly identified single-host PyVs (Fig 3), 19 PyVs shared 68-83% nt sequence identities with the closest-known species or with each other in the LTAg coding sequences, therefore were considered to be novel PyV species according to a demarcation standard based on ≥ 15% genetic divergence in the LTAg gene [36]. Seven PyVs shared 86-90% identity with the closest known species or with each other in the LTAg gene, therefore showing insufficient divergence; however, all were identified for the first time in their specific bat hosts. Furthermore, these PyVs were exclusively detected in bat species separated by considerable distances (≥ 564 ± 328 km) (Fig 3 and S2 Table). They could therefore be considered novel PyV species, a proposal further reinforced by additional correlation provided by the genetic selection analysis described below [36]. Ra-Hl PyV and Ra-Mh PyV, exhibited 66% and 77% nt sequence identities in their LTAg coding sequences with the closest known PyV, could therefore be proposed as novel PyV species. However, the nomenclature for these PyVs requires further discussion as inter-family host-switching is unprecedented for mammalian PyVs [36].

### Phylogenetic analysis and identification of host-switching events

In order to verify the host-switching events, an LTAg amino acid tree was constructed to compare the 28 new PyVs with 126 known *Alpha*- and *Betapolyomavirus* species (including 38 bat-borne PyV reference species). As shown in Fig 4, 28 new bat PyVs (colored in the figure) clustered within the *Alphapolyomavirus* (n=19) and *Betapolyomavirus* (n=9) genera along with 38 known bat PyVs. Within each viral genus, bat PyV sequences grouped into 7 clusters, referred to here as Alpha 1-7 and Beta 1-7, respectively. Among these novel PyVs, two viruses, Ra-Hl PyV in the *Alphapolyomavirus* and Ra-Mh PyV in the *Betapolyomavirus* genera, were identified in bat hosts of different families living in the same colony, suggestive of two viral host-switching events, labeled as host-switching event 1 and 2 (HS1 and 2 in Fig 4). Furthermore, a third PyV clustered in the phylogenetic tree with PyVs carried by bats of different families (Hipposideridae and Rhinolophidae), indicative of another inter-family host-switching event (HS3 in Fig 4).

**Fig 4.**
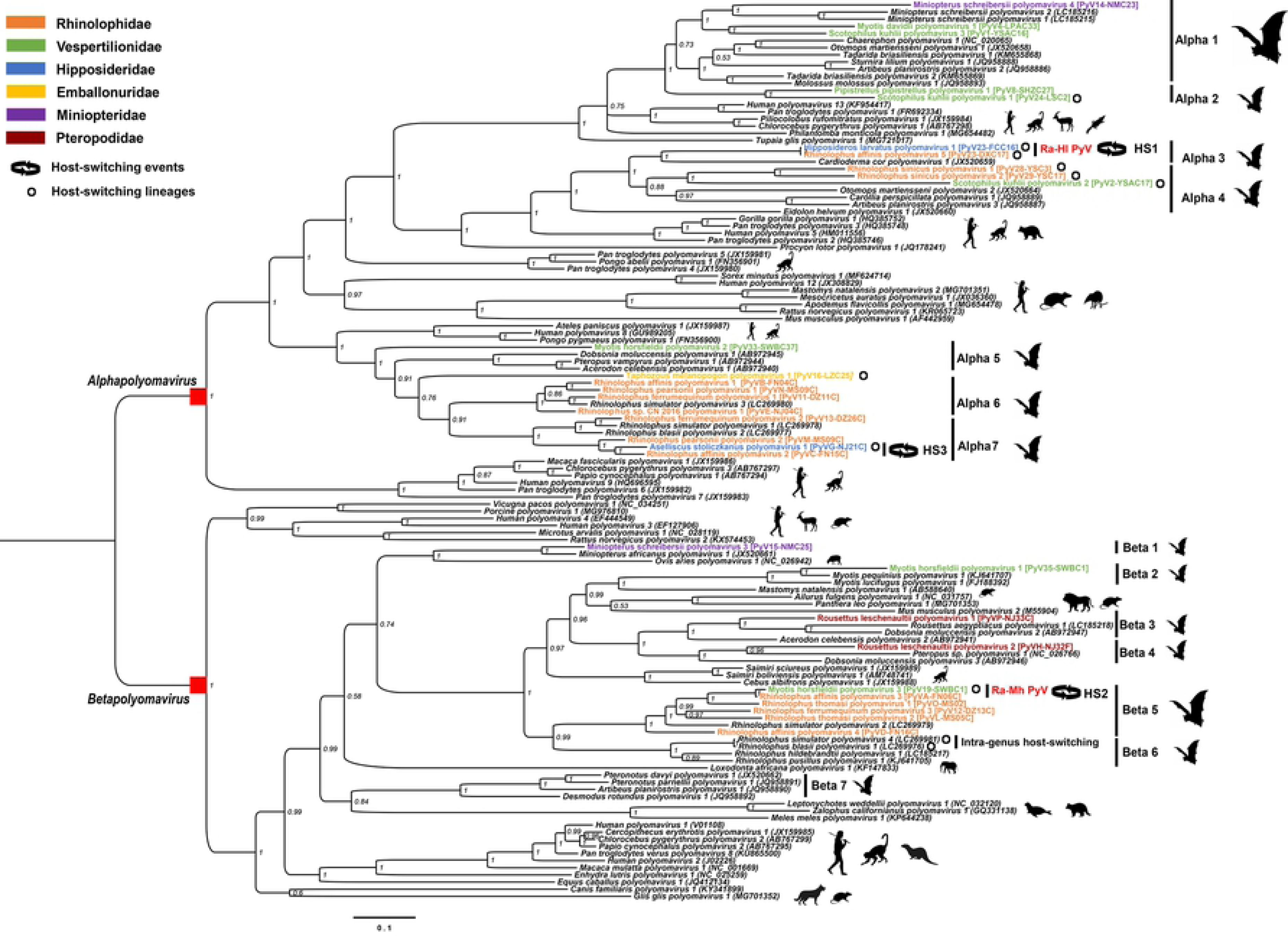
Bayesian phylogenetic tree of the PyV LTAg protein sequences. The tree was constructed with 156 amino acid sequences of 28 new PyVs identified in the present study, and 126 PyV references (including 38 bat PyV species references) in *Alpha-* and *Betapolyomavirus* genera available in GenBank. Names of new PyVs with their strain names in parentheses at the branch tips are colored differently according to their host species within the Chiropteran family, as shown in the top left-hand corner. Three host-switching events are marked as HS1 to HS3 by opposing arrows. Host switching lineages identified by correlation analyses are also shown with black circles. Silhouettes of the mammalian hosts of PyVs are shown alongside each clade. Bayesian posterior probability supporting the branching is shown next to the branches. Abbreviations of PyVs: Ra-Hl PyV, Rhinolophus affinis-Hipposideros larvatus polyomavirus; Ra-Mh PyV, Rhinolophus affinis-Myotis horsfieldii polyomavirus.

The first host-switching event HS1 in the Alpha 3 cluster (Fig 4) occurred between *Rh. affinis* (family Rhinolophidae) and *H. larvatus* (family Hipposideridae). Strikingly, Ra-Hl PyV, characterized in *Rh. affinis* (Rhinolophus affinis polyomavirus 5) and *H. larvatus* (Hipposideros larvatus polyomavirus 1), exhibited 100% nt identity between both genomes. Importantly, they were detected in multiple animals and, also, from different colonies. The Rhinolophus affinis polyomavirus 5 was detected in two colonies (colonies 16 and 17 separated 10 km apart in south Guangxi province in southern China) with detection rates of 1/22 and 8/28, respectively. Notably, the Hipposideros larvatus polyomavirus 1 was also detected in colonies 16 and 17 in *H. larvatus*, also in Guangxi province, with positive rates of 3/19 and 2/10, respectively.

The host-switching event HS2 in the Beta 5 cluster (Fig 4) occurred between *Rh. affinis* (family Rhinolophidae) and *My. horsfieldii* (family Vespertilionidae), involving Ra-Mh PyV characterized in *Rh. affinis* (Rhinolophus affinis polyomavirus 3) and in *My. horsfieldii* (Myotis horsfieldii polyomavirus 3). These PyV strains were also extremely close genetically but not identical, sharing 97-99% nt genomic identity. Ra-Mh PyV was frequently detected in both hosts: Rhinolophus affinis polyomavirus 3 was detected in five colonies of *Rh. affinis* bats in Yunnan, Guangxi and Fujian provinces (23%; 43/188), in which 7 full genomes shared 97-100% nt identity in their LTAg coding sequences (marked with black circles in Fig 3). The Myotis horsfieldii polyomavirus 3 had a high positive rate 63.8% (23/36) in *My. horsfieldii* bats within a single colony (colony 24 Fig 1 and S1 Table) where, notably, *Rh. affinis* and *My. horsfieldii* species roosted together. In this cave, *Rh. affinis* also harbored Rhinolophus affinis polyomavirus 3 with a 57% (12/21) detection rate. Geographically, Rhinolophus affinis polyomavirus 3 was widely distributed in *Rh. affinis* hosts.

The third host-switching event, HS3, in the Alpha 7 cluster (Fig 4) occurred between *A. stoliczkanus* (family Hipposideridae) and *Rhinolophus* spp. (family Rhinolophidae). The Aselliscus stoliczkanus polyomavirus 1 exhibited 87% and 86% nt identity with Rhinolophus affinis polyomavirus 2 and Rhinolophus pearsonii polyomavirus 2, respectively (Fig 3). The two sampling locations of the *Rhinolophus* species were from caves located 731 km and 86 km respectively from the cave in which the single Aselliscus stoliczkanus polyomavirus 1 positive sample was detected.

Besides these three host-switching events, a PyV-positive rectal tissue sample, named Pipistrellus pipistrellus polyomavirus XJPp02 [strain PyV10-YXAC25], collected from a *Pipistrellus pipistrellus* (family Vespertilionidae) bat in colony 28 in northwestern Xinjiang exhibited 98% nt identity in a partial VP1 fragment (∼270 nt) with Rattus norvegicus polyomavirus 1 (detected in German rats) [35] in the *Alphapolyomavirus* genus (marked as host-switching event HS4 in Fig 2). A qPCR protocol designed to detect both the previously described rat PyV and the *P. pipistrellus* PyV was employed to screen samples comprising spleen, kidney, lung, liver, rectal and brain tissues in all *P. pipistrellus* bats (n=158) collected in three locations in Xinjiang in 2016. Also screened were rectal and kidney tissues from brown rats (n=51) available from collections in two other locations in Xinjiang during 2015-2016 [38]. The original PyV10-YXAC25 isolate yielded CT values of 36-38 upon repeated testing, indicative of very low copy numbers of target. An additional lung tissue sample derived from another *P. pipistrellus* bat from the same colony in which the original PyV10 was detected was also positive, with a higher viral load (CT: 34) by qPCR. No rat samples were positive. Despite repeated attempts to amplify by inverse PCR, no genomes were recovered from the bat samples.

### Correlation of genetic divergences between PyVs and bat hosts

As shown in Fig 3 and 4, although seven new PyVs showed high LTAg nt identities (86-90%) with each other or the closest known PyVs, the phylogenetic clustering supported codivergence events rather than host-switching events. For instance, Rhinolophus pearsonii polyomavirus 1 and Rhinolophus affinis polyomavirus 1 shared 88% nt identity (Alpha 6); Rhinolophus affinis polyomavirus 1 and Rhinolophus sp. CN 2016 polyomavirus 1 shared 85% identity (Alpha 6); Rhinolophus ferrumequinum polyomavirus 2 shared 89% identity with Rhinolophus simulator polyomavirus 1 (LC269978) previously identified in Zambian bats (Alpha 7); and Rhinolophus ferrumequinum polyomavirus 1 shared 90% identity with Rhinolophus simulator polyomavirus 3 (LC269980) also identified in Zambian bats (Alpha 6). These last two instances showed that PyVs from Chinese and African horseshoe bats were remarkably similar despite the geographical separation.

To investigate further the factors potentially influencing the divergence between PyVs with high nt sequence identities, Mantel’s correlation tests were performed to compare the divergence among PyV genome sequences with those among *cytb* genes of their corresponding hosts and to control the putative effects of the geographic location on the bat collection. As shown in Table 3, correlation analysis was performed using three groups of data: 1) a total of 42 PyVs with full genomes from the present study; 2) a total of 48 PyVs including the 42 PyV genomes described here and 6 PyV genomes we previously identified in Zambian horseshoe bats [26]; 3) a total of 37 PyVs obtained by removing 11 PyVs (marked in Fig 4 which potentially arose by host-switching events) from the 48 PyVs identified in both China and Zambia. Analysis of each of these three groups of data showed that there was negligible correlation between geographical distances and divergence for either hosts or viruses. Moreover, the geographical proximity between the colonies had little effect on the co-evolution of viruses with their respective hosts.

**Table 3.**
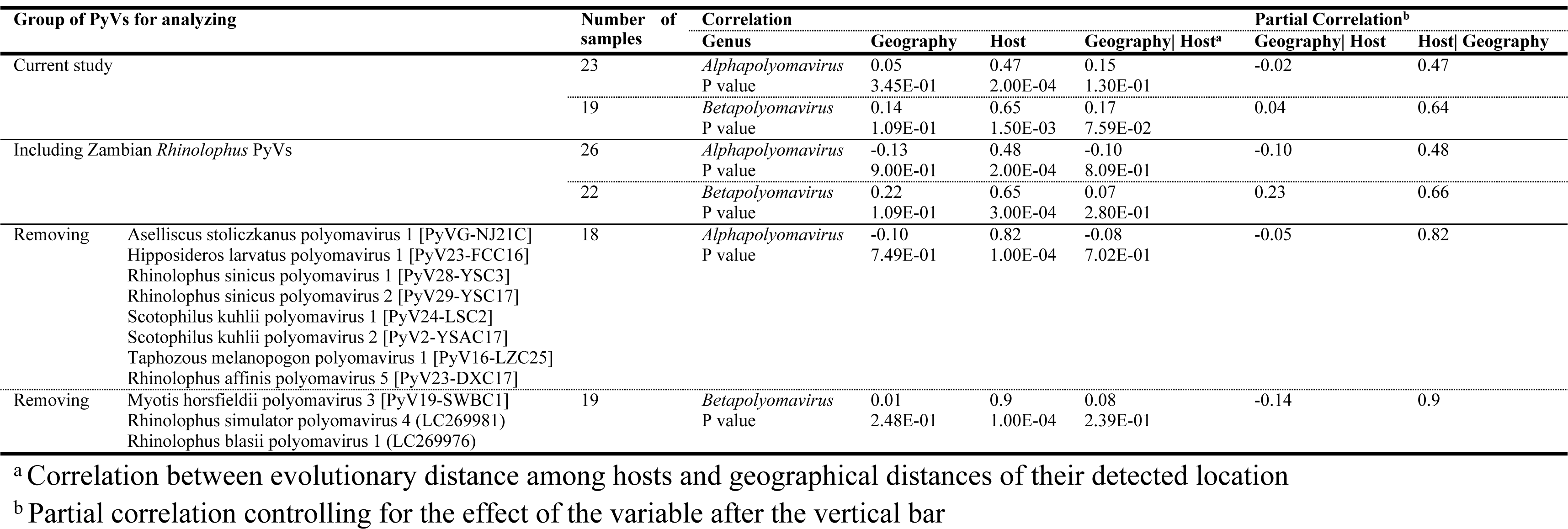
Correlation analyses of genetic divergences between PyVs and bat hosts.

Analysis using the first group of data showed that a significant correlation existed between host and PyV divergence per genus considering the 42 genomes identified in China. The correlation between bat hosts and PyVs in the *Alpha-* and *Betapolyomavirus* genera was 0.47 (*P* < 2×10^-4^) and 0.65 (*P* < 1.5×10^-3^), respectively. This first dataset supported the intra-host divergence model of PyV and hosts, with multiple viral lineage duplications, in support of a previous model of PyV evolution [17]. The addition of PyV sequences sampled from African *Rhinolophus* spp. [26] increased the correlation index between host and viral divergence as the number of considered sequences increased. Altogether, these findings support an intra-host divergence model for the radiation of PyV species within the order Chiroptera. In the third group, by removing 8 sequences (also marked as host switching lineages in Fig 4) in the *Alpha-* and 3 in the *Betapolyomavirus* genera, which, notably, included host-switching events (HS1-HS3) from both PyV genera, the correlation indices increased 40% and 30%, respectively.

### Bayesian estimation of tMRCA of bat hosts and PyVs

To test whether the host-switching events of the PyV species occurred after the divergence of host species, the times to most recent common ancestor (tMRCA) of both bat hosts and the PyVs were estimated employing a Bayesian approach. As shown in Fig 5A, the tMRCA of bat hosts were estimated based on the *cytb* sequences (∼1800 bp) of bats that harbored the 28 novel PyVs. The results showed that divergence among *Rhinolophus* species happened 14.2 to 17.3 million years ago (MYA) with a 95% highest posterior density (HPD), which is consistent with previously reported divergence times (16–17 MYA) of the bat hosts based on mitochondrial DNA [39] and other sampled bat families [40]. The Bayesian results for PyV divergence times (Fig 5B) also showed that the tMRCAs for both *Alpha*- and *Betapolyomavirus* genera were 115-182 MYA with 95% HPD, overlapping with the divergence times of the tMRCA of placental mammals (92 to 116 MYA) [41]. However, when compared with the Bayesian results for the bat hosts, the time scale of PyV divergence was not always in agreement: specifically, 1) the divergence time between *Rhinolophus* and *Hipposideros* hosts was estimated with a median 36.3 MYA in the 95% HPD range (34, 38) MYA, while the tMRCAs of Rhinolophus affinis polyomavirus 5 and Hipposideros larvatus polyomavirus 1 were estimated with a median of 0.02 MYA in the 95% HPD range (0.0012, 0.12) MYA; 2) The tMRCAs of Myotis horsefieldii polyomavirus 3 and Rhinolophus affinis polyomavirus 3 had a median of 1.05 MYA in the 95% HPD range (0.4, 1.9) MYA, while the median tMRCA of both corresponding host species was 66.76 MYA in the 95% HPD range (60.9, 73.5) MYA; 3) The median tMRCA of Rhinolophus affinis polyomavirus 2 and Aselliscus stoliczkanus polyomavirus 1 was estimated as 11.61 MYA in the 95% HPD range (6.6, 17.8) MYA, however, the tMRCA between both hosts had a median of 36.3 MYA in the 95% HPD range (34.3, 38.2) MYA. Finally, 4) the estimated time of divergence between Scotophillus kuhlii polyomavirus 2 and Pipistrellus pipistrellus polyomavirus 1, with a median of 37 MYA in the 95% HPD range (23, 54) MYA, was less than the predicted divergence time between the hosts, with a median of 44.42 MYA in the 95% HPD range (40, 49) MYA. In the four examples, the most parsimonious explanation for the lesser median ages to divergence of the PyVs compared to the longer tMRCAs for the bat hosts is that these represent examples of inter-family viral host-switching events.

**Fig 5.**
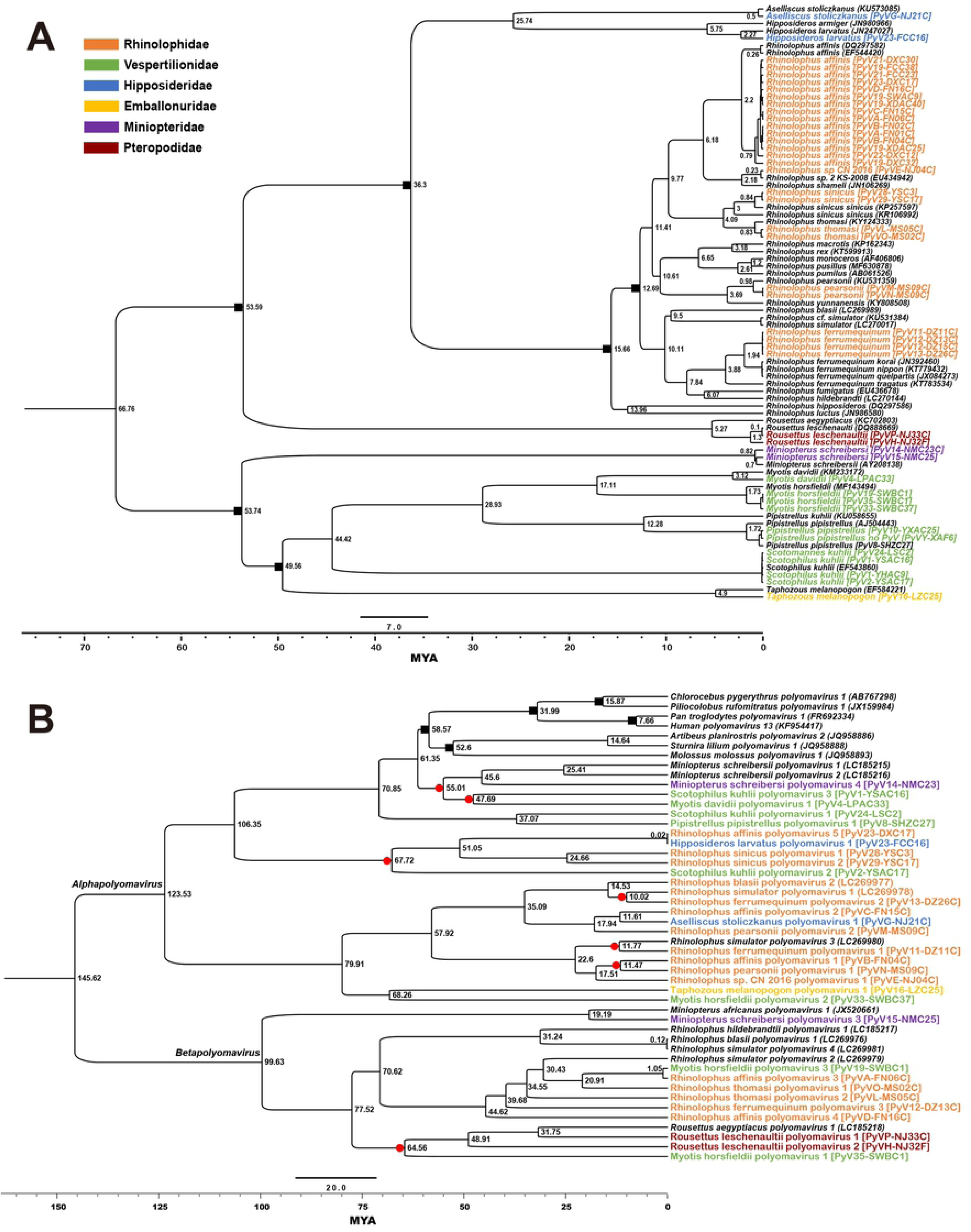
Dating the tMRCA of host bat species employing mitochondrial *cytb* sequences (A) and PyV species using the LTAg gene (B). *Cytb* and PyV sequences from this study are colored according to the Chiropteran family of the host species and following the legend in the top right-hand corner. Calibration points of *cytb* are marked on the branches with black squares (A). Calibration points using PyVs identified from non-human primates and other bat species are shown in the upper section of the phylogeny and marked with black squares (B). The timescale at the bottom of both figures and the tMRCAs shown next to the branches are in millions of years (MYA) before the present. Median age of the tMRCAs of *cytb* (A) and PyVs (B) are shown next to the branches. In Fig 5B, branches that matched the divergence between host species in Fig 5A are identified with red circles.

### Intra-lineage host duplication and host-switching events during bat PyV evolution

The findings of the Mantel test and other studies [17] supporting intra-host divergence for PyVs, required reconciliation with the compelling evidence of similar viral species infecting evolutionarily distant bat species. Co-phylogenetic reconstruction analyses using JANE 4 were therefore performed separately for both viral genera. The best-supported solution to explain the divergence in the genus *Alphapolyomavirus* for the novel Chinese PyVs and African samples was modeled with 11 co-speciations, 4 lineage duplications, 13 lineage losses and 9 host-switching events (Fig 6A). Among the 9 inferred host-switching events, in agreement with the tMRCA estimations, it was implied that Rhinolophus affinis polyomavirus 5 originated in the lineage which gave rise to Hipposideros larvatus polyomavirus 1, Rhinolophus sinicus polyomavirus 1 and 2, and Scotophilus kuhli polyomavirus 2 indicating that Rh-Hl PyV was originally a PyV harbored by *Rh. affinis*, which was subsequently transmitted to *H. larvatus*. The best-supported solution also suggested that the lineage of Rhinolophus affinis polyomavirus 2 had given rise to Aselliscus stoliczkanus polyomavirus 1. Other host-switching events inferred earlier in the divergence among PyVs were more difficult to corroborate with the available tMRCA data. Divergence in the genus *Betapolyomavirus* involved 6 co-speciations, 2 lineage duplications, 7 lineage losses and 7 host-switching events (Fig 6B). Similar to the *Alphapolyomavirus* genus, the solution for the *Betapolyomavirus* genus suggested, among the 7 host-switching events, that the origin of Myotis horsfieldii polyomavirus 3 was as a descendant of the lineage giving rise to Rhinolophus affinis polyomavirus 3 (i.e. that Rh-Mh PyV was originally a PyV harbored by *Rh. affinis* and transmitted to *My. horsfieldii*), in agreement with the tMRCA results. In both solutions, although the presence of duplication events could explain the existence of more than one different viral species in the same host species, in multiple instances the dissimilarity between viral species from the same host species was more easily explained by host-switching events following the divergence between host species.

**Fig 6.**
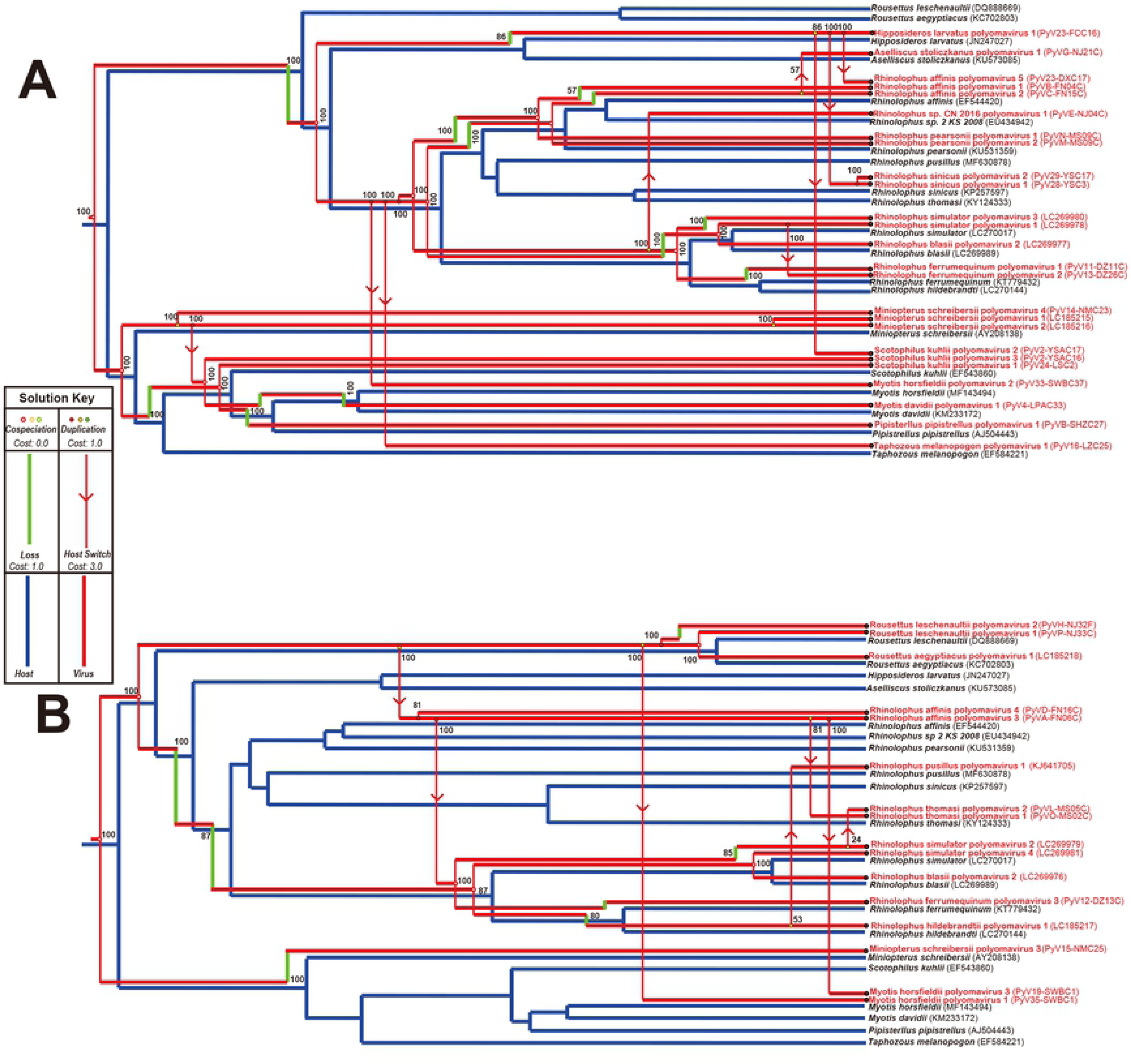
Co-phylogenetic models for bat PyVs in *Alpha-* and *Betapolyomavirus* genera with their hosts. The blue phylogenies show the inferred host divergence. The red phylogenies are annotated to reconstruct the divergence among viral species according to JANE 4 in the *Alphapolyomavirus* (A) and *Betapolyomavirus* (B) genera. Symbols for the annotations are shown in the solution key legend and percentage support for the events are shown next to their annotations.

### Positive selection in divergence of Chinese and African *Rhinolophus* PyVs are focused on the external surface of the VP1

Given the diversity of viral species identified in the genus *Rhinolophus*, viral proteins under positive selection during the evolution of the different PyV lineages were of interest as possible speciation determinants. We performed a Bayesian analysis to calculate the ω rate (dN/dS) to identify genomic regions of the *Alpha*- and *Betapolyomavirus* genera in *Rhinolophus* PyV clusters, which were likely subject to positive selection (S3 Fig). As expected, the majority of the horseshoe bat PyV genomes were found to be under neutral or negative selection (ω values ≤ 1); however, a number of residues in VP2, VP1, LTAg and STAg proteins did show evidence of positive selection (ω values > 1) in both viral genera (shown in red in S3 Fig) and amino acid residues relative to human BK virus (species of *Human polyomavirus 1*) and simian virus 40 (species of *Macaca mulatta polyomavirus 1*) are detailed in S5 Table. We focused on the major capsid protein VP1 as this determines antigenicity and the engagement of sialylated glycans present on host glycoproteins and glycolipids during cellular attachment and entry in mammalian (murine, human and monkey) viruses [42-46].

In the *Rhinolophus* alphapolyomavirus cluster, 11 residues were found under positive selection, located in four regions of VP1 (S5 Table): in the BC-loop (D77, N79 and V82), the DE-loop (L135, A139 and T143), the HI loop (N274, N275 and S279) and the C-terminal residues I367 and N369. In the *Rhinolophus* betapolyomavirus cluster, 9 residues had ω values > 1 at the N-terminus (K17), in the BC loop (A75, A77, T80, V82 and P84), the DE loop (N134 and Q149) and the EF loop (S178). No positively selected amino acids were identified in the interior of the major capsid protein.

## Discussion

While mammalian PyVs are highly host specific [12] there are data suggesting cross-species jumps of PyVs between closely related mammalian hosts. Previously, short-range intra-genus host-switching events of PyVs were identified in African horseshoe bats, occurring between *Rh. simulator* and *Rh. blasii* in a single Zambian cave [26]. Strikingly, the genomes of PyVs characterized in respective bat species were 99.9% identical and contained only four virus species-specific polymorphisms. Importantly, these polymorphisms were found to be present in both spleen and kidney samples of each bat host species, indicating a systemic and productive infection. The genetic stability and restricted host-specificity of mammalian PyVs had previously suggested a model of strict codivergence for PyV evolution [47]. However, this is inconsistent with family-wide patterns of diversity, where an important contribution of intra-host lineage duplication, recombination and, possibly, host-switching events is evident. A model of intra-host divergence, with these additional evolutionary processes, occurring over the ancient timescales for association of PyVs with multicellular life, has been proposed to explain the extant diversity within the family *Polyomaviridae* [17]. The findings in the present study extend these observations to PyVs in Chinese horseshoe bats and other bat hosts (see model in Fig 7) and provide the first definitive evidence of inter-family host-switching of PyVs in mammals occurring among the bat families Rhinolophidae, Hipposideridae and Vespertilionidae.

**Fig 7.**
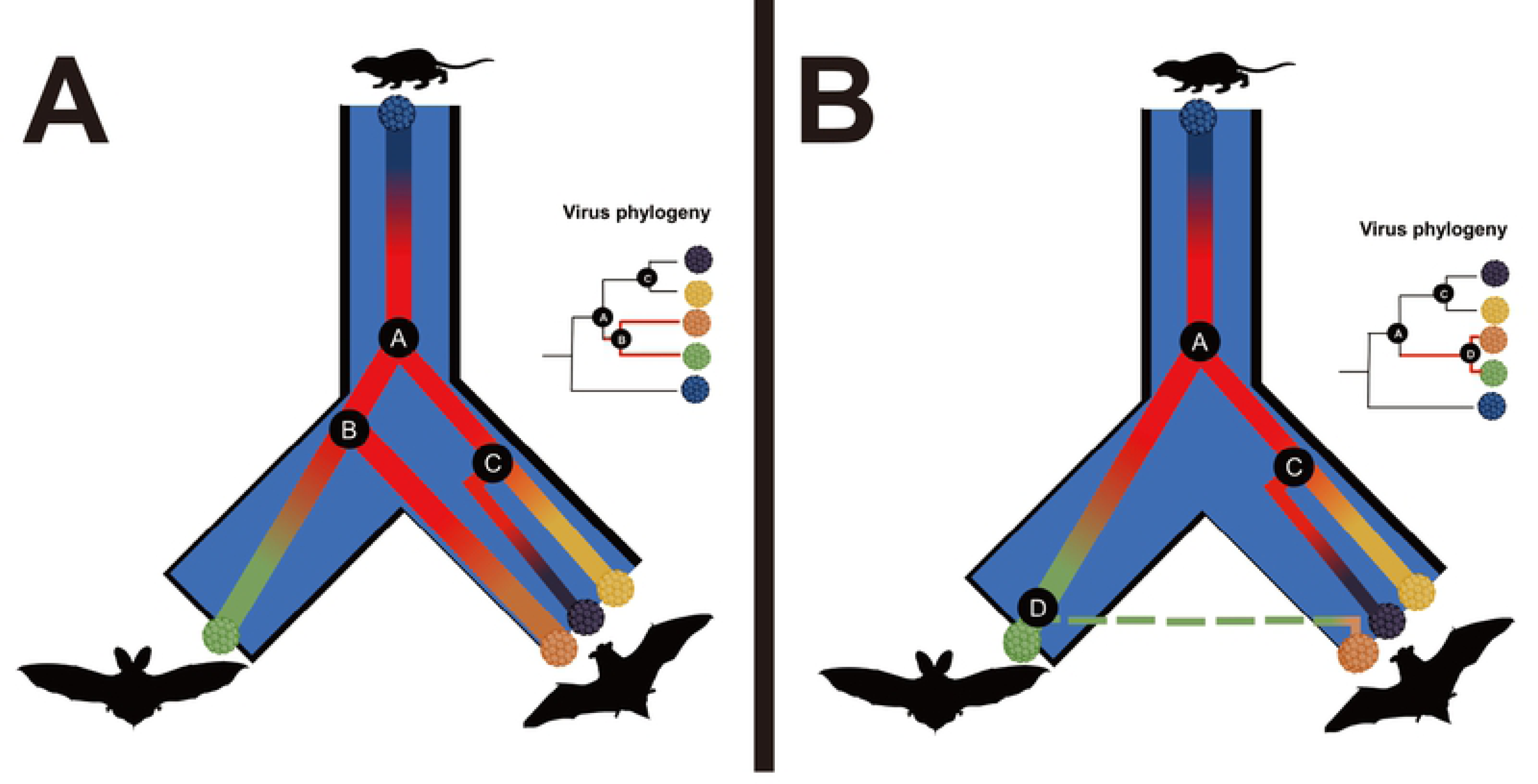
Model of PyV evolution in mammals by intra-host divergence with infrequent host-switching events. The panel illustrates the models for: A) intra-host divergence and B) intra-host divergence with infrequent host-switching events. In both panels, the bifurcating fork in blue represents the divergence from a common ancestor of two mammalian host species, with the overlapping co-divergence of viral species in color. Divergence events are marked A-D representing: (A) viral duplication before the divergence of host species, (B) co-divergence with the host species, (C) viral duplication in the same host species and (D) duplication and host-switching to a diverged (susceptible) host species. The phylogenies close to the cartoons are the expected phylogenetic relationship for each respective case with the events marked at the nodes.

The host-switching events of PyVs identified in the present study indicate that transmission of PyVs between different bat species has been an ongoing process during the evolutionary history of PyVs. HS1 and HS2 occurred between bats of different families and strains of PyVs sharing 100% or 97-100% nt identity were characterized in their respective hosts. The tMRCA analysis estimated that HS1 and HS2 occurred between 0.02-1.05 MYA. Considering the “slow” evolutionary rate of PyVs (8 × 10^-9^ substitutions per site per year as previously estimated [48]), the transmission events of Rh-Hl PyV and Rh-Mh PyV in bats happened recently, with a small number of mutations accumulating in the viral genomes during viral adaptation to the permissive hosts. Compared with HS1 and HS2, HS3 was estimated to have occurred much earlier, between 6.6-17.8 MYA, with the Aselliscus stoliczkanus polyomavirus 1 showing 87% nt identity with the closest PyV (Rhinolophus affinis polyomavirus 2), indicating that the former had been transmitted from Rhinolophus bats to A. stoliczkanus bats for a long period of time, had co-evolved with A. stoliczkanus and, finally, had formed a novel PyV species. In addition to these observed host switching events, the tMRCA estimation and correlation analysis showed strikingly that numerous PyVs had also been derived from host switching, thereby shaping the evolutionary history of PyVs and providing evidence that inter-species transmission of PyVs, at least in bats, is not as rare as previously thought.

An issue raised by the present study is how to define and name a PyV species when inter-family host-switching events of PyV are identified. The ICTV criteria for the formal designation of PyV species in mammals must include the binomial name of the host species [36]; nevertheless, the inter-family host-switching events of PyVs discovered in the present study as well as the inter-species host-switching event also identified in *Rhinolophus* hosts in a previous study [26] which has challenged this nomenclature system for bat PyVs. Furthermore, numerous host-switching lineages have also been characterized here by correlation analyses and dating estimation. All these findings suggest that host-switching of PyVs has occurred not only in different bat species but also at different time points in different geographical locations. The ICTV nomenclature criteria should therefore be updated to include recognition of PyVs derived from two different mammalian host species.

Inter-species transmission has already been observed in bird PyVs such as budgerigar fledgling disease PyV (BFDV), which can infect multiple avian species with pronounced pathogenicity [49]. Nevertheless, the ICTV nomenclature has utilized only the order level of the host species in the naming scheme: for instance, species belonging to BFDV have been classified as *Aves polyomavirus 1*. This naming scheme could also be expanded using the more descriptive host species, as with the mammalian PyV species. So far, PyVs with host-switching events have only involved two host species, illustrating the limited ability of PyVs for transmission between different mammalian hosts. Therefore, the improved naming scheme of novel PyVs with host-switching events proposed here would include the binomial names of both host species; e.g., Rhinolophus affinis-Hipposideros larvatus polyomavirus (Rh-Hl PyV), identified in the present study, would become the prototype of a new species named *Rhinolophus affinis-Hipposideros larvatus polyomavirus 1*.

Another ICTV criterion is the allowable genetic distance of the LTAg coding sequence (≥ 15%) to demarcate PyV species. This cut-off value is not strict, as exemplified by two PyVs identified in two different species of squirrel monkeys which shared 89% sequence identity but were recognized as two separated PyV species [36]. In the present study, we have discovered numerous novel PyVs carried by diverse bat species, all exhibiting 86-90% sequence identities with their closest known species or with each other. These should be considered new PyV species since all were detected exclusively in distinct bat species in different areas and exhibiting strict host specificity. Examples include the PyVs identified in different *Rhinolophus* bat species from China and Africa sharing 89-90% nt identify. The correlation and genetic selection analysis reported here has shown that the high sequence identity of different PyVs carried by different host species was due to the co-divergence with strong negative selection of PyVs in respective hosts, indicating that the ≥ 15% cut-off in LTAg gene identity for species demarcation is perhaps no longer adequate and should be updated to ≥10%. With regard to the relationship between Chinese and African bat PyVs referred to above, it is of interest to note that hepadnaviruses (pararetroviruses with DNA genomes) identified in Chinese *Rhinolophus* bats in another study [49] were also found to exhibit the closest genetic relation (79.6–79.7% nt identify) with viruses carried by African *Rhinolophus* bats, [50] thereby suggesting that co-divergence of DNA viruses within their *Rhinolophus* bat hosts is relatively common.

In the present study we have identified 28 novel PyVs from 15 bat species in 6 families [36], a finding that has considerably increased our knowledge of both the genomic diversity and the evolution of PyVs. Additionally, since there is a dearth of commercially available, standardized bat cell lines for isolation and identification of bat-borne viruses there are now added options to establish immortalized bat cell lines from many bat species by using LTAg genes cloned from divergent PyVs identified in specific Chinese bat species. At present, the Tb1 Lu cell line derived from the free-tailed bat (*Tardarida brasiliensis*, family Molossidae) is currently the only bat cell line available in the American Type Culture Collection (ATCC Nr. CCL-88) [51]. Recently, however, Banerjee *et al*. [52] have reported immortalization of primary bat kidney cell lines from the big brown bat (*Eptesicus fuscus*, family Vespertilionidae) which were stably transformed by a plasmid harboring a little brown bat (*Myotis lucifugus*, family Vespertilionidae) PyV LTAg. The ability to establish cell lines from different tissues of diverse bat species and to make these available to researchers is a prerequisite for replication studies, virus-host interactions and new virus discoveries.

Papillomaviruses (PVs), which are closely related to PyVs and share similar properties such as, high genetic stability and strict host specificity, have been shown to cause transient and potentially cross-species infections. García-Pérez *et al*. [53] reported that PVs do not follow strict host restrictions in free-ranging Iberian bats (*Eptesicus serotinus*, family Vespertilionidae), as they found evidence of host-switching events between two different bat species (*E. serotinus* and *E. isabellinus*). They therefore recommended a revised paradigm of strict host specificity in PVs [53].

The findings presented here have possible implications for the study of zoonotic transmissions involving highly virulent viral pathogens in bats. For example, horseshoe bats within the family Rhinolophidae have been found to harbor genetically diverse severe acute respiratory syndrome (SARS)-like corona viruses, which acted as major reservoirs for spillover of SARS coronavirus to humans [7]. The Middle Eastern respiratory syndrome (MERS) coronavirus has similarly been associated with severe disease and lethal respiratory infections, mostly in Saudi Arabia. Likely transmitted via dromedary camels, it has been shown to be closely related to several bat coronaviruses, including those sequenced from species within the family Vespertilionidae (*Neoromicia capensis, Pipistrellus abramus and Vespertilios uperans*) [54]. Recently, swine acute diarrhea syndrome (SADS) coronavirus has been associated with high morbidity and mortality in pig populations in Guangdong province, China in close proximity to the origin of the SARS pandemic, and striking similarities between the SADS and SARS outbreaks in geographical, temporal, ecological and etiological settings exist [9]. We consider noteworthy that in the present study we identified host-switching events of the normally highly host-restricted PyVs within the bat families Rhinolophidae and Vespertilionidae. Whether this reflects a greater propensity for host-switching in these bat families when compared with others and what mechanisms are involved require further functional studies. Conceivably, given the high prevalence we and others have detected of bat PyVs, co-infection of bat species within the families Rhinolophidae and Vespertilionidae may affect the prevalence and/or transmission of CoVs and other high virulent pathogens in bats. This suggestion warrants further study.

Our findings that inter-family transmission of PyVs can occur in mammals may also have implications for studies of viral oncogenesis proposed previously for the *Polyomaviridae*. For example, three bovine PyVs (BoPyVs) have been identified in cattle [55] with more likely given the 13 so far identified as infectious agents in humans [56]. Intriguingly, BoPyV2, occupies the same clade as Merkel cell PyV which is causally implicated in oncogenic transformation in a neuroendocrine carcinoma of the elderly. BoPyV3, is also related to human polyomavirus 6 which has been reported in non-melanoma skin cancers, specifically keratocanthomas [57]. Colorectal cancer has been suggested to have an infectious disease etiology [58] and the relatively high thermoresistance of PyVs may preclude inactivation by lowered cooking temperatures [59]. Similarly, in field studies, sequences of BK virus were identified by metagenomics in the feces of non-human primates [60]. Our evidence that inter-family transmission of PyVs can occur in bats provides a precedent for studies of longer-range transmission of PyVs in mammals.

In addition to three bat inter-family host-switching events of PyVs, we also identified a partial VP1 fragment in rectal tissue of a *P. pipistrellus* bat collected in northwestern Xinjiang that showed 98% nt identity with the VP1 of a previously described Rattus norvegicus polyomavirus 1 identified in Germany [35]. Further qPCR assay also detected the virus in lung tissue of another *P. pipistrellus* bat but not from any brown rat (n=51) collected in the same Xinjiang area. This preliminary observation provides an impetus to investigate potential PyV host-switching between mammal species beyond families.

The host-switching events identified in this study were strongly associated with the ecological environment of the hosts, with host-switching only identified between different bat species in one to two colonies. Our study suggests that PyVs can transmit between different mammalian hosts between two different animal families by frequent contact of hosts within specific environments, such as the densely packed bat populations that occur in large hibernacula in caves. The host-switching PyVs were identified from rectal tissues of multiple individuals, indicating that shedding via feces is a key factor for viral transmission by the fecal-oral route between different host species.

PyV attachment to target cells is mediated by cell surface receptors. MHC class I acts as a co-receptor and SV40 employs the ganglioside GM1 as receptor [61]. The finding that gangliosides are also primary receptors for human PyVs via VP1 [62] suggests that this may be a general property for primate PyVs although whether this is applicable to other mammalian PyVs, including bats, is unknown. In the present study, amino acid residues in VP1 were found to be under positive selection in *Rhinolophus* PyVs in the BC, DE and HI loops (*Alphapolyomavirus*) and in the BC, DE and EF loops (*Betapolyomavirus*). Interestingly, both *Rhinolophus* PyV genera showed evidence of positive selection in the external surface BC and DE loops. In BK virus, serological subtypes are assigned based on mutations present within the BC loop [63]. It has also been previously demonstrated that sialylated glycans can act as primary receptors for PyVs [64] and binding to sialic acid moieties was attributed to amino acids residues within the BC, DE and HI loops, suggesting that these VP1 domains play a key role in virus attachment and cellular entry events [45]. The surface loops of VP1, i.e. the BC, DE, HI and EF loops, are responsible for viral antigenicity and escape mutants cluster in these regions, suggesting they represent conformational epitopes recognized by host neutralizing antibodies [65, 66]. VP1 surface loops form a unique virus-host interaction conformation that determines the host range, cellular tropism and pathogenicity which may explain the enrichment of positive selected residues within these regions [67]. The availability of annotated bat genomes in the future, as proposed by the genome consortium Bat1K [68], and annotation of the genes encoding putative entry receptors (such as, for example, the cell surface ganglioside and other sialylated glycan receptors) from within the bat families Rhinolophidae, Hipposideridae and Vespertilionidae from species previously found to harbor identical or closely related viruses would facilitate the functional characterization of the cellular receptors/attachment factors and of which amino acid residues in VP1 are required for binding and entry. Availability of annotated bat genomes would also help determine whether certain families of bats are more susceptible than others to virus-host switching events.

## Materials and methods

### Ethics statement

The procedures for sampling bats and brown rats in this study were reviewed and approved by the Administrative Committee on Animal Welfare of the Institute of Military Veterinary Medicine (Laboratory Animal Care and Use Committee Authorization permits JSY-DW-2010-02 and JSY-DW-2016-02). All live animals were maintained and handled according to the Principles and Guidelines for Laboratory Animal Medicine (2006), Ministry of Science and Technology, China.

### Sample information

The tissues of 1,083 bats in the present study were archived and sub-packed samples were stored at −80°C following collection between 2015-2016 from 29 colonies distributed across the provinces of Yunnan, Fujian and Zhejiang, Guangxi and Xinjiang in China. Archived samples of rectal and kidney tissues from brown rats (n=51) were collected in Xinjiang between 2015-2016 [38]. Detailed information of sampled bats and rats is shown in S1 Table.

### Polyomavirus screening and complete genome sequencing

Approximately 50 mg samples of rectal and lung tissues from the 208 bats in colonies 1-4 collected in Yunnan province were pooled and subjected to viral metagenomic analysis, as per our previously published method [34]. Due to the complexity of the PyV-related reads detected by the viral metagenomic analyses, the PyV-specific nested PCR protocol [69] based on the VP1 gene developed by Johne *et al*. in 2005 [27] was employed to screen all metagenomic samples, and partial VP1 amplicons were cloned and bidirectionally sequenced by standard methods. In addition, the PCR was expanded to screen genomic DNA extracts from rectal tissues of bats collected from other Chinese provinces.

Genomic DNA from each bat tissue sample was extracted manually using the TIANamp Genomic DNA Kit (Tiangen), according to the manufacturer’s protocol. PCR screening employed the PCR master mix (Tiangen) with the following thermocycling conditions: 45 cycles for both outer and inner PCRs with denaturation at 94°C for 30 s, annealing at 46°C (outer PCR) or 56°C (inner PCR) for 30 s or 1 min, and extension at 72°C for 1 min, with ddH2O as a negative control. Positive PCR amplicons were purified by gel extraction (Axygen) and sequenced directly on a 3730xl DNA Analyzer (Genewiz) to confirm homology to PyVs and then ligated into vector pMD-18T (TaKaRa) and used to transform *E. coli* DH5α competent cells (Tiangen). Four clones from each VP1 amplicon were picked for Sanger sequencing. GenBank accession numbers of 192 partial PyV VP1 sequences are showed in S3 Table.

Oligonucleotide primers for inverse PCRs for amplification of the 5 kb PyV genomes were design based on the partial VP1 sequences. Multiple pairs of hemi-nested degenerate primers were subsequently designed for VP1 clusters which showed close phylogenetic relationships (see S6 Table). All ∼5 kb products were amplified from bat genomic DNA by inverse PCR employing the PrimeSTAR MAX DNA polymerase (Takara), purified by gel extraction (Axygen), cloned into pCR-Blunt II-TOPO vector (Invitrogen) and bidirectionally sequenced by primer walking to Phred quality scores > 30 by 3730xl DNA Analyzer (Genewiz). After removal of vector sequences, the full genomes of each PyV were assembled by Geneious software using the partial VP1 and 5 kb products.

A real-time PCR (qPCR) method (S6 Table) was established based on a 59 nt fragment of the VP1 genes of Pipistrellus pipistrellus polyomavirus XJPp02 and Rattus norvegicus polyomavirus 1 was designed to screen spleen, kidney, lung, liver, rectal and brain tissues in all the *P. pipistrellus* bats (n=158) collected in three locations in Xinjiang in 2016 and also rectal and kidney tissues from brown rats (n=51) available from collections in two other locations in Xinjiang during 2015-2016 (S1 Table). The extracted DNAs from the tissues were amplified using the Probe qPCR kit (TaKaRa) as per the manufacturer’s protocol on a Stratagene Mx3000P.

### Phylogenetic analysis

The pairwise identity of the LTAg coding sequences was calculated with Sequence Demarcation Tool (SDT) v1.2 [70]. The nucleotide sequence of host *cytb*, the sequences of PyV genes and proteins were aligned using MAFFT under the algorithm FFT-NS-i [71]. Phylogenetic trees were inferred using MrBayes v3.2, with 10^6^ generations chains to ensure < 0.01 standard deviations between the split frequencies and 25% of generations discarded as burn-in [72], as described previously [73]. For the LTAg multiple-sequence alignment, a mixed substitution model was used to explore the best substitution model when inferring the topology. On the other hand, the best fitting nucleotide substitution models for host *cytb* and the PyV LTAg coding sequence alignments were chosen with MEGA7 [74] by choosing the lowest Bayesian information criterion (BIC).

The tMRCA for hosts and viral samples were estimated with Bayesian analyses conducted as 4 independent MCMC chains of 10 million generations each, sampled every 1,000 generations (BEAST v2.4.6 [75]). The *cytb* and LTAg gene multiple sequence alignments were used to infer the divergence times among hosts and PyVs, respectively. The general time-reversible substitution model with gamma-distributed rate variation across sites and a proportion of invariable sites (GTR+G+I) were assumed for both models. Other settings were: uncorrelated lognormal relaxed molecular clocks for host and PyV models with a Bayesian skyline population and exponential population models, respectively. These models chosen had the highest likelihood when compared to other models with the PathSampler application in BEAST v2.4.6 package. The host model was calibrated with divergence times between genera as inferred by Agnarsson [40] and the PyV model was calibrated following Buck *et al*. [17, 76].

### Correlation analysis

The divergence correlation between the hosts and PyVs in the *Alpha-* and *Betapolyomavirus* genera was assessed by applying Mantel’s correlation tests to the pairwise evolutionary distance matrices for hosts and viral samples in both genera with 10^4^ repetitions. Additionally, the matrices were also analysed against the geographical distance between sampling points, and partial correlations between host and polyomavirus distance matrices were estimated controlling for the correlation with the geographical distance. The correlation analysis was performed considering three sets of data per genus: 1) considering only samples identified in China, 2) considering samples identified in China and samples identified in Africa [26], and 3) removing samples from 2) such as the correlation index increased. Removed samples in 3) were chosen by estimating the correlation coefficient without each sample and then removing samples whose absence maximized the correlation between hosts and PyVs.

### Co-phylogenetic analysis

Mapping the divergence of PyV species to the divergence among hosts was performed with JANE 4 [77]. The solutions for *Alpha-* and *Betapolyomavirus* genera were generated with costs for co-speciation, duplication, loss and host switch events as 0, 1, 1 and 3, respectively. The input was the phylogenetic tree for the host species inferred with mitochondrial *cytb* sequences, the phylogenetic trees for the *Alpha-* and *Betapolyomavirus* genera inferred with amino acid sequences of the LTAg protein and the table indicating the association between hosts and viruses.

### Analysis of natural selection on the evolution of proteins in *Rhinolophus* PyV species

The ω ratio (dN/dS) was estimated for each coding region in clades of *Rhinolophus* PyV species in *Alpha-* and *Betapolyomavirus* genera with MrBayes v3.2.6 [72] with NY98 as the codon model for 5×10^4^ generations sampling every 250 and assuring the effective sample size was > 100 for estimated parameters. Tridimensional structure of the VP1 protein was modeled by homology with SWISS-MODEL [78].

## Supporting information

**S1 Fig**. **Maximum-likelihood phylogenetic tree of partial VP1 nucleotide sequences from PyV positive samples**. The tree was constructed with 192 partial VP1 sequences of PyVs in present study (filled circles), and 74 PyV references (including 38 bat PyV species references) in *Alpha-* and *Betapolyomavirus* genera downloaded from GenBank. These 192 partial VP1 sequences were classified into 44 clusters representing host-switching events (green circles; clusters 7, 10, 17 and 31; red circles: no host switching). GenBank accession numbers of 192 partial PyV VP1 sequences in this tree are listed in S3 Table. The tree was generated in MEGA7 by the maximum-likelihood method based on the TN93 model and evaluated with 250 bootstrap replicates.

**S2 Fig**. **Bayesian phylogenetic tree of the *cytb* sequences of bats**. The tree was constructed with 37 reference bat *cytb* genes and 42 bat *cytb* genes in the present study (colored differently according to the bat families) for confirmation of the species of bats from which PyV full genomes were obtained. The posterior probability supporting the branching is shown next to the branch. GenBank accession numbers of *cytb* sequences in this tree are listed in S4 Table.

**S3 Fig**. **Analysis of positive selection in PyVs infecting the horseshoe bat *Rhinolophus* in *Alpha-* and *Betapolyomavirus* genera**. The top panels show the ω rate (dN/dS) estimated with a Bayesian approach for clusters of *Rhinolophus* PyVs in both *Alpha-* and *Betapolyomavirus* genera with positions adjusted to Rhinolophus affinis polyomavirus 1 [PyVB-FN04C] and Rhinolophus affinis polyomavirus 3 [PyV19-SWAC9], respectively. Tridimensional models for the VP1 proteins corresponding to both genera were inferred by structure homology modelling. The top and bottom views of both modelled proteins are shown in gray with sites under positive selection colored in red.

**S1 Table**. **Summary of sampled colonies and detailed information of PyV positive bats following a geographic approach of the collection sites**.

**S2 Table**. **Summary of detected PyVs and their positive rates in colonies**.

**S3 Table**. **GenBank accession numbers of partial PyV VP1 sequences**.

**S4 Table**. **GenBank accession numbers of 1**.**8 kb *cytb* sequences**.

**S5 Table**. **VP1 amino acid residues under positive selection in Rhinolophus affinis polyomavirus 1 in the *Alphapolyomavirus* genus and Rhinolophus affinis polyomavirus 3 in the *Betapolyomavirus* genus with homologous positions relative to human BK virus (species of *Human polyomavirus 1*) and Simian virus 40 (species of *Macaca mulatta polyomavirus 1*)**.

**S6 Table**. **List of oligonucleotides used in the present study**.

